# A meta-analysis of the effect of organic and mineral fertilizers on soil microbial diversity

**DOI:** 10.1101/2020.10.04.325373

**Authors:** Daniel P. Bebber, Victoria R. Richards

**Author notes:** Author for correspondence:* Daniel P. Bebber.

## Abstract

Organic agriculture, employing manures or composts, has been proposed as a way of mitigating undesirable impacts of mineral fertilizer use. Of particular interest is the effect of fertilizer regime on soil microbes, which are key to nutrient cycling, plant health and soil structure. However, the effect of fertilizers on soil microbial diversity remains poorly understood. Since biological diversity is an important determinant of ecosystem function and a fundamental metric in community ecology, the effects of fertilizer regimes on soil microbial diversity are of theoretical and applied interest. Here, we conduct a meta-analysis of 37 studies reporting microbial diversity metrics in mineral fertilized (NPK), organically fertilized (ORG) and unfertilized control (CON) soils. Of these studies, 32 reported taxonomic diversity derived from sequencing, gradient gel electrophoresis, or RFLP. Functional diversity, derived from Biolog Ecoplate™ measures of carbon substrate metabolism, was reported in 8 studies, with 3 studies reporting both diversity metrics. Bacterial and archaeal diversity was reported in 28 taxonomic studies, and fungal diversity in 8 taxonomic studies. We found that functional diversity was 2.8 % greater in NPK compared with CON, 7.0 % greater in ORG vs CON, and 3.8 % greater in ORG vs NPK. Bacterial and archaeal taxonomic diversity was not significantly different between NPK and CON, but on average 2.9% greater in ORG vs CON, and 2.4 % greater in ORG vs. NPK. Fungal taxonomic diversity was not significantly different between any treatment pairs. There was very high residual heterogeneity in all meta-analyses of soil diversity, suggesting that a large amount of further research is required to fully understand the influence of fertilizer regimes on microbial diversity and ecosystem function.

## INTRODUCTION

Diversity plays a key role in the resilience and adaptability of complex systems (Page, 2011), and biological diversity has been central to understanding of the structure and function of ecological communities (Ricklefs and Schluter, 1994). Human activities are rapidly eroding global biodiversity (Ceballos et al., 2015), hence understanding how human activities influence diversity and how negative impacts can be avoided is an important goal of applied ecology (Rudd et al., 2011; Sutherland et al., 2006). Recent advances in environmental DNA sequencing and metabarcoding have revealed enormous and unexpected microbial diversity in all habitats, particularly soils (Lloyd et al., 2018; Thompson et al., 2017). Given the fundamental importance of soils in terrestrial ecosystems, agriculture and food security, there is growing interest in the role of microbial diversity in processes such as nutrient cycling (Delgado-Baquerizo et al., 2016), and how soil diversity can be managed for maintenance of ecosystem services (Lemanceau et al., 2015). Soil health, the capacity of soils to function as a living system and sustain and promote plant and animal communities, is linked to microbial diversity (Kibblewhite et al., 2008; van Bruggen et al., 2019). Microbial diversity is correlated with soil ecosystem multifunctionality including plant productivity, microbial biomass, availability of nitrate, ammonium and phosphorus, and nitrogen mineralisation rates (Delgado-Baquerizo et al., 2016). In agriculture, soil microbes are critical to desirable functions such as nutrient cycling, carbon storage, erosion control via soil aggregation, and disease suppression (Mazzola, 2004; Rillig et al., 2002; Sahu et al., 2017). A large fraction of the world’s agricultural soils are in poor and deteriorating condition largely due to agricultural activities (Wuepper et al., 2020). Therefore, understanding how to manage soils for microbial diversity could help to prevent further deterioration.

Modern intensive, or conventional, agricultural methods, including application of mineral and chemical fertilisers, regular tillage, and use of synthetic pesticides and herbicides, aim to increase soil nutrition and suppress harmful species to produce higher crop yields, but these methods are environmentally damaging (Reganold and Wachter, 2016). For example, growing pressure on agricultural land has resulted in soil nutrient depletion and soil erosion (Wuepper et al., 2020). Mineral fertilizers supply nitrogen (N) as ammonium nitrate or urea, phosphorus (P) and potassium (K), with around 100 Tg N applied globally each year (FAO, 2020). Typically less than half the N applied is taken up by crops, the remainder contributing to water pollution and the release of NOX greenhouse gases (Zhang et al., 2015). Organic agriculture, which applies manure- or compost-based fertilizers and soil conditioners and does not employ chemical pest controls (with certain exceptions), has been proposed as a means of both increasing soil nutrition and reducing environmental impacts (Luo et al., 2018; Reganold and Wachter, 2016), but evidence for organic benefits remains equivocal. For example, eutrophication potential appears greater in many organic than conventional crop systems (Clark and Tilman, 2017)

Soils are exceedingly complex and biologically-diverse ecosystems, varying in physico-chemical and biological composition across spatial scales (Howe et al., 2014; Thompson et al., 2017). Meta-analyses of multiple individual studies can reveal general trends and patterns in this complexity and determine the effects of management interventions. For example, meta-analysis of observational data suggest that biological richness (measured by DNA sequences) is greatest at neutral pH and at a mean temperature of 10 °C (Thompson et al., 2017). Meta-analysis of experimental studies suggests that pH has the greatest influence on soil microbial diversity among global change factors, alpha diversity rising with pH; that nitrogen and NPK have negative or non-significant effects on alpha diversity depending on microbial group, with significant negative influences of N alone on diversity in agricultural soils; that soil functionality, defined as the range of biogeochemical processes carried out by soils, increases with N and NPK; and that changes in diversity are negatively correlated with changes in functionality, perhaps due to functional redundancy (Zhou et al., 2020). Other meta-analyses have demonstrated that microbial diversity (Venter et al., 2016) and biomass (McDaniel et al., 2014) increase in crop rotations compared with monocultures. A meta-analysis of experimental studies showed that organic agriculture greatly increases microbial biomass carbon, microbial biomass nitrogen, and enzymatic activity compared with conventional systems (Lori et al., 2017). Another meta-analysis found that organic amendments increase crop yields by supporting soil microbial activity, but this did not investigate effects on soil microbial diversity (Luo et al., 2018). A small meta-analysis found no significant effect of organic agriculture on soil organism diversity, but this only included five studies and occurred before widespread use of sequencing to soil microbial diversity (Bengtsson et al., 2005). However, despite growing application of sequencing technologies in microbial ecology, the effects of organic and mineral fertilizers on measures of soil microbial diversity remain unsynthesized.

Here, we conduct a meta-analysis quantifying the effects of organic fertilizers (manures and manure composts) and mineral fertilizers (NPK) on soil microbial diversity, in comparison with unfertilized controls. We compare results for taxonomic and functional diversity, and investigate the influence of factors such as soil chemistry and duration of organic treatment on these effects. We hypothesise that organic inputs will increase microbial diversity as compared to mineral fertilizer inputs, based on the observation that organic fertilizers increase microbial biomass and enzymatic activity compared with NPK (Lori et al., 2017). Ecological theory and observational data suggest that soil microbial diversity has a hump-shaped relationship with microbial biomass, with fungal and bacterial diversity increasing with biomass for all but the highest biomass levels (Bastida et al., 2021).

## MATERIALS AND METHODS

### Criteria for meta-analysis

We used meta-analysis to test for differences in soil microbial functional (H_fun_) and taxonomic (H_tax_) diversity between soils treated with manure-based organic fertilizer (ORG), mineral fertilizer (NPK) and control (CON). Here, H_tax_ is defined as Shannon’s diversity index calculated from the relative abundance of species, operational taxonomic units (OTUs) or amplicon sequence variants (ASVs; Callahan et al., 2017) in a sample. Shannon’s diversity index was reported in all valid studies, hence we do not consider any other diversity indices (e.g. Simpson’s) that were reported additionally in a small number of studies (e.g. Ma et al., 2018). H_fun_ is defined as Shannon’s diversity calculated from the relative conversion rates of various carbon sources in Biolog Ecoplates (e.g. Ros et al., 2006). We conducted a literature search using combinations of the terms ‘soil’, ‘organic’, ‘agriculture’, ‘microbial’, ‘diversity’, ‘bacteria*’, ‘fung*’, ‘communit*’, ‘fertilizer’, ‘manure’ and ‘compost’ (where * indicates a wildcard search, where appropriate) on Google Scholar, Scopus and Web of Science between June 2021 and October 2021. Additionally, the reference lists of the papers were browsed to find potentially appropriate studies which were not identified during the online literature search. All potentially appropriate papers identified for the meta-analysis came from peer-reviewed journals.

Papers were considered eligible for the analysis if they met the following criteria, adapted from Lori et al. (2017): Comparisons of the farming systems should be pairwise, meaning the organic- and mineral-fertilised treatments were subject to the same climatic conditions before and during sampling; The organic treatment must have been applied for a minimum of two consecutive years prior to soil sampling, and the treatments must be defined by the study; Results must report the mean Shannon diversity index (H) per treatment, uncertainty of the mean (either standard error of the mean or standard deviation of the sample distribution) and sample size (n); experiments were conducted in open fields or under cover, not in pots or containers; the study must have been published no earlier than the year 2000 because of the lack of comparable methods before this date; the mineral and organic treatments must be clearly described (description of fertilizer, application rate). Non-manure organic fertilizers were not included (e.g. Wang et al., 2016). We only included experimental studies in which the only variable was fertilizer treatment, and NPK and organic treatments were applied. If there were several different organic or conventional treatments within the same study, all appropriate combinations were reported and treated as individual comparisons but with non-independent errors (see meta-analysis methods). Results of combination treatments (e.g. manure plus NPK fertilizer) were not included (e.g. Zhao et al., 2014).

For all valid studies, we extracted mean, SE or SD, and sample size for H, for control, NPK and organic treatments. Where SE was reported, this was converted to SD by multiplying by the square root of sample size. When data were presented in graphical form, we extracted means and errors using the online tool WebPlotDigitalizer (https://automeris.io/WebPlotDigitizer/). Additionally we abstracted the following variables: study location; reported soil type, duration of organic treatment, organic treatment type (i.e. source of manure), organic treatment application rate (mass per hectare per year or equivalent nitrogen mass per hectare per year); mineral fertilizer application rate (nitrogen mass per hectare per year); crops grown; taxonomic group of microbes analysed (e.g. all, bacteria, fungi); median soil depth sampled (usually the mid-point of the range of depths sampled); brief summary of the methodology used (e.g. 16S sequencing, 16S DGGE, Biolog Ecoplate). Where reported, we abstracted mean and SE of soil pH, and mean and SE of available soil nitrogen content (mg kg^-1^) which was often reported as the concentration of ammonium and nitrate.

### Meta-analysis

Meta-analysis was conducted using the *metafor* package v. 2.4–0 for R v. 4.0.2 (Viechtbauer, 2010). Effect size was calculated as the log–transformed ratio of means, log(*y*_exp_/*y*_con_) where *y*_exp_ and *y*_con_ are the means of the experimental treatment and control, respectively (Hedges et al., 1999). There was at least one pairwise comparison of H in each study. However, several studies reported data from more than one pairwise comparison, with multiple organic and/or conventional fertilisation treatments investigated. In addition, several studies were conducted at the same research site. We fitted multi-level random– and mixed–effects (i.e. testing for the effects of moderator variables) models by restricted maximum likelihood, with random intercepts per location. Other similar meta-analyses have ignored potential non-independence among results from the same study or location and treated each reported mean as an independent replicate (e.g. Lori et al., 2017). Standard errors and confidence limits for parameters were estimated from the *t*–distribution. We report the *I*^2^ statistic and *Q*-test for heterogeneity (Higgins and Thompson, 2002), and provide funnel plots rank correlation tests for funnel plot asymmetry (Viechtbauer, 2010).

## RESULTS

We abstracted data from 37 studies which met our requirements (Table S1). Our search criteria provided a long-list of 267 studies, of which 199 were rejected on the basis of the abstract, leaving a shortlist of 68. A further 31 were rejected after detailed reading of the full text. Reasons for the rejection of studies included: no reporting of diversity metrics (e.g. reporting of species richness); no or unclear reporting of standard error or standard deviation for diversity metrics; no manure used in organic treatment; no organic treatment alone (e.g. manure only applied in combination with NPK); no NPK treatment; no experimental treatments (e.g. comparison of different farms under organic and conventional treatments); lack of detail in description of treatment types or levels.

Around half (21) of the included studies reported one set of comparisons, with either or both of unfertilized control (CON) and mineral fertilizer (NPK), 10 reported two comparisons, and five studies reported up to six comparisons, giving 65 comparisons in total. The majority of treatments were replicated three or four times. The functional diversity studies, and a single study reporting soil dilution plate assays (Mahanta et al., 2017), did not differentiate between fungi, bacteria and archaea. The single soil dilution plate assay was omitted from the meta-analysis due to the low sensitivity of the method and is not considered further here. In that study, poultry manure treatment resulted in significantly greater microbial diversity than inorganic fertilizer treatment (Mahanta et al., 2017). In the taxonomic diversity studies, bacteria were the most commonly analysed group (26 studies, including one of actinomycetes only, one of bacteria and archaea, and one of nitrogen-fixing bacteria), followed by fungi (8 studies, including one of arbuscular mycorrhizal fungi only). Most studies were conducted in China (24), followed by India (6), with single studies from Austria, Canada, Denmark, Kenya, Korea, the Netherlands, and the USA.

Soil types were not reported according to any standard taxonomy. Soils were sampled to a median depth of up to 15 cm (median 10.0 cm, IQR 7.5 - 10.0 cm). Soil nitrogen content in unfertilized plots was reported in 16 studies (mean mg N kg^-1^ median 36.3, IQR 27.7 – 86.3). Manure-based organic fertilizers were applied in all studies, derived variously from cattle, pigs, poultry, horses, sheep, or mixtures of these. Composted manures were employed in 9 studies, fresh manures in 26 studies, and one study employed a mix. A small number of studies reported more than one manure treatment or level (X. Hu et al., 2018; Liu et al., 2019; Mahanta et al., 2017). Application rates were reported as either total mass per area per year (30 studies, median 16.8 t ha^-1^ y^-1^, IQR 5.1 – 19.6 t ha^-1^ y^-1^) and/or as nitrogen addition (18 studies, median 103.3 kg ha^-1^ y^-1^, IQR 75.0 – 223.8 kg ha^-1^ y^-1^). We did not attempt to estimate nitrogen content in manures where this was not reported, due to the large vari ability in the nitrogen fraction among different treatments. The reported nitrogen fraction varied from 0.4 % in horse manure to 2.7 % in pig manure, with overall median 1.0 % (IQR 0.7 – 2.0 %) among the treatments reported in the studies. The level of nitrogen in the manure treatment was similar to that in corresponding NPK fertilizer treatments, where reported (absolute percentage difference median 0.1 %, IQR 0.0 – 29.9 %). The duration of organic treatment was right-skewed, but many studies reported treatment periods of several decades (median 22 y, IQR 13 – 34 y). A variety of crops were grown, either as monocultures or in mixtures, the most common being maize (12 studies), wheat (10 studies), and rice (6 studies). Soil pH in unfertilized plots was reported in 18 studies (mean pH median 6.4, interquartile range 5.9 - 7.5). Meta-regression showed that soil pH declined under NPK treatment compared to control soil (Fig. S1a). Manure-based fertilizers increased pH in acid soils and reduced pH in alkaline soils (Fig. S1b). Funnel plots showed no asymmetry (Fig. S2). There was a large degree of residual heterogeneity in the meta-analyses of soil pH (Table S2).

H_fun_ was reported by 8 studies, all employing Biolog Ecoplates (Table S1). H_fun_ varied between 1.12 and 4.61 with values in three clusters comprising a single study from Austria with H_fun_ ∼ 1.2, four studies from China and one from India with H_fun_ ∼ 2.9, and two studies from India with H_fun_ ∼ 4.5 (Fig. 1a-c). H_fun_ increased by an average of 2.8 % in NPK vs. CON, 7.0 % in ORG vs. CON, and 3.8 % in ORG vs. NPK (Fig. 2, Table 1). We found no influence of the duration of organic amendment on the change in H_fun_ compared with control (LRM change per year of organic treatment -0.0008 ± 0.0013 y^-1^, t = -0.58, p = 0.56) or NPK (0.0006 ± 0.0007 y^-1^, t = 0.84, p = 0.40) treatments. We found no effect of the level of NPK addition relative to control (−0.0013 ± 0.0009, t = -1.42, p = 0.15), nor of level of organic fertilizer addition quantity (0.0000 ± 0.0000, t = -0.35, p = 0.72) or nitrogen equivalent relative to control (0.0001 ± 0.0005, t = 0.18, p = 0.86). We found marginally-significant effect of control soil pH on the effect size of ORG vs. CON (−0.057 ± 0.018, t = -3.2, df = 2, p = 0.088), but not of soil pH on NPK vs. CON (0.0055 ± 0.0432, t = 0.128, df = 2, p = 0.91). Funnel plots did not exhibit significant asymmetry (Fig. S3). There was a large degree of residual heterogeneity in all meta-analyses of H_fun_ (I^2^ > 75 %, Table S2).

**Table 1.** Summary statistics for meta-analysis of functional diversity (H_fun_) differences between fertilizer treatments and control. One study (Ros et al., 2006) reported three comparisons, hence a multi-level random model was employed to control for within-study effects. Studies (N) refers to the number of studies and number of comparisons. LRM is the log-transformed ratio of means ± SE and 95 % Confidence Interval. Difference (%) is the effect-size back-transformed to percentage difference.

| Comparison | Studies (N) | LRM | LRM (95% CI) | Difference (%) | Difference (95% CI) |
| --- | --- | --- | --- | --- | --- |
| NPK vs. CON | 6 (8) | 0.027 ± 0.004 | 0.018, 0.037 | 2.8 | 2.0, 3.6 |
| ORG vs. CON | 7 (9) | 0.068 ± 0.012 | 0.041, 0.095 | 7.0 | 4.6, 9.5 |
| ORG vs. NPK | 7 (9) | 0.037 ± 0.014 | 0.004, 0.070 | 3.8 | 0.9, 6.7 |

**Fig. 1.**
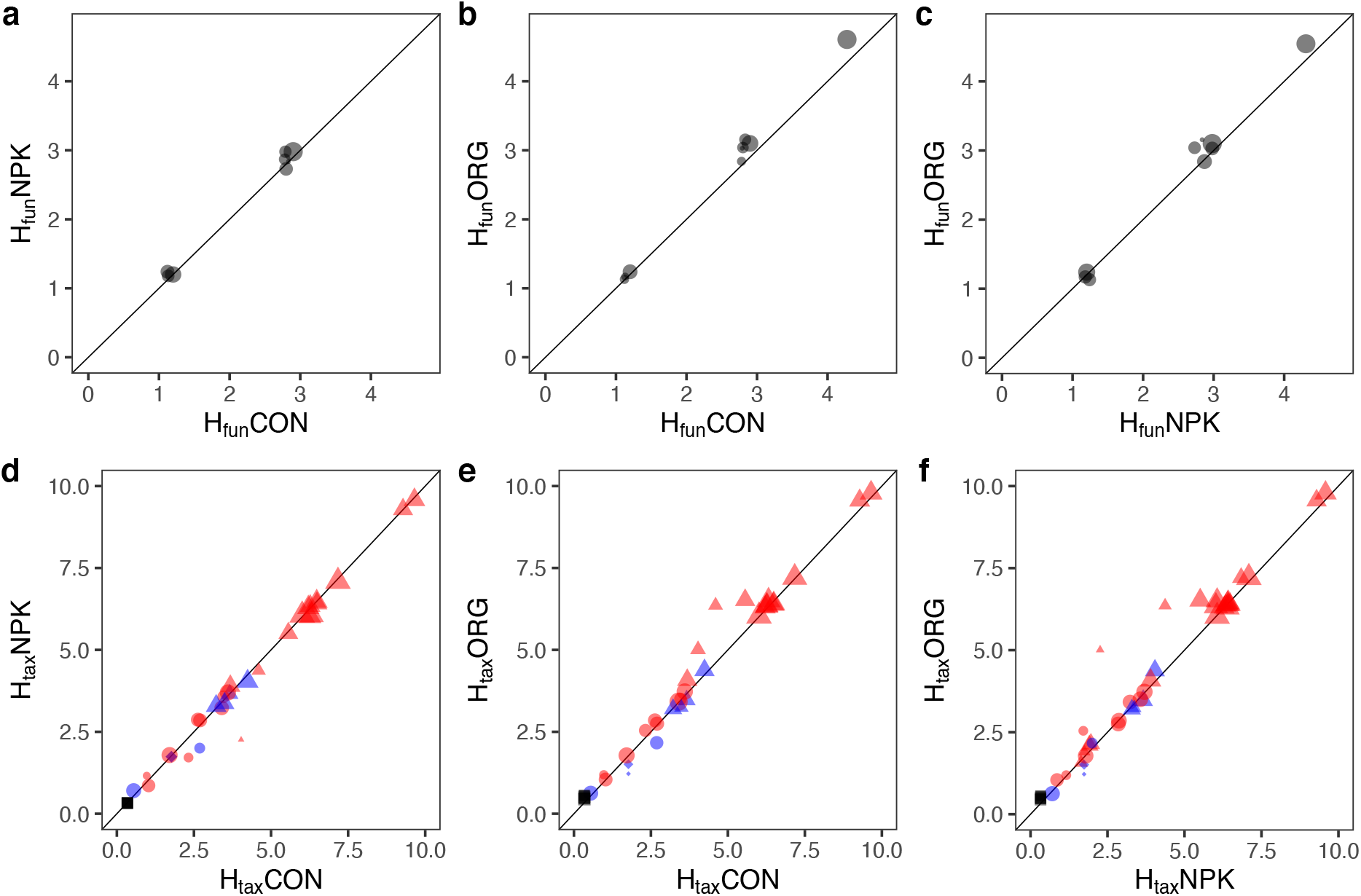
Comparison of H_fun_ (a-c) and H_tax_ (d-f) in (a,d) NPK vs. CON, (b,e) ORG vs. CON, (c,f) ORG vs. NPK. Results for fungi are in blue, prokaryotes in red. Points show reported values with size proportional to log(1/variance). Gel electrophoresis results are circles, sequencing results are triangles, TRFLP are diamonds.

**Fig. 2.**
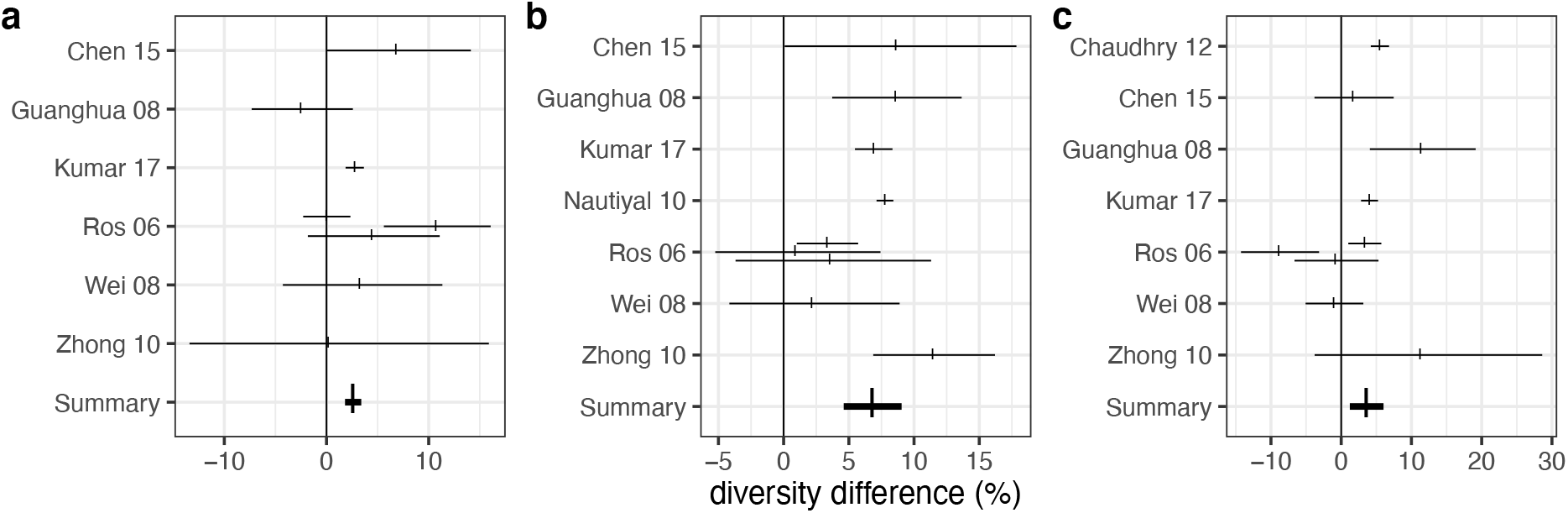
Functional soil diversity differences. a) NPK vs. CON, b) ORG vs. CON, c) ORG vs. NPK. Horizontal bars and ticks show 95% confidence intervals and means for effect sizes from the log-transformed ratio of means, back-transformed to give percentage differences. Summary estimates are given at the bottom of each plot. Random intercepts were fitted per Study.

H_tax_ was reported in 31 studies, employing DNA amplicon sequencing (21 studies), various forms of gradient gel electrophoresis (9 studies), and T-RFLP (1 study). H_tax_ tended to be higher in bacteria and archaea (hereafter, prokaryotes) than fungi, and in sequencing compared with DGGE methods (Fig. 1d-f). Median H_tax_ for prokaryotes was 6.40 using sequencing and 2.63 using DGGE, in controls. Median H_tax_ for fungi was 3.95 using sequencing and 1.62 using DGGE, in controls. H_tax_ of NPK-fertilized soils did not differ from control for prokaryotes or fungi (Fig. 3, Table 2). H_tax_ of prokaryotes was significantly greater in ORG soils than both NPK and CON (Fig. 3, Table 2). There was no significant difference in fungal H_tax_ between ORG and CON or NPK (Fig. 3, Table 2). Funnel plots did not exhibit significant asymmetry (Fig. S4). There was a large degree of residual heterogeneity in meta-analyses of H_tax_ (I^2^ > 95 %, Table S2).

**Table 2.** Summary statistics for meta-analysis of taxonomic diversity. Statistics as in Table 1.

| Comparison | Class | Studies<br>(N) | LRM | LRM (95%<br>CI) | Change<br>(%) | Change % (95<br>% CI) |
| --- | --- | --- | --- | --- | --- | --- |
| NPK vs.<br>CON | Fungi | 8 (10) | $0.026 \pm 0.016$ | -0.007, 0.058 | 2.6 | -0.6, 5.9 |
| | Prokaryotes | 22 (29) | $-0.011 \pm 0.014$ | -0.017, 0.040 | 1.1 | -1.7, 4.0 |
| ORG vs.<br>CON | Fungi | 8 (10) | $0.028 \pm 0.014$ | -0.002, 0.055 | 2.7 | -0.2, 5.7 |
| | Prokaryotes | 22 (29) | $0.029 \pm 0.008$ | 0.011, 0.047 | 2.9 | 1.1, 4.8 |
| ORG vs.<br>NPK | Fungi | 8 (10) | $-0.017 \pm 0.015$ | -0.048, 0.014 | -4.2 | -4.7, 1.4 |
| | Prokaryotes | 27 (39) | $0.024 \pm 0.011$ | 0.001, 0.046 | 2.4 | 0.1, 4.7 |

**Fig. 3.**
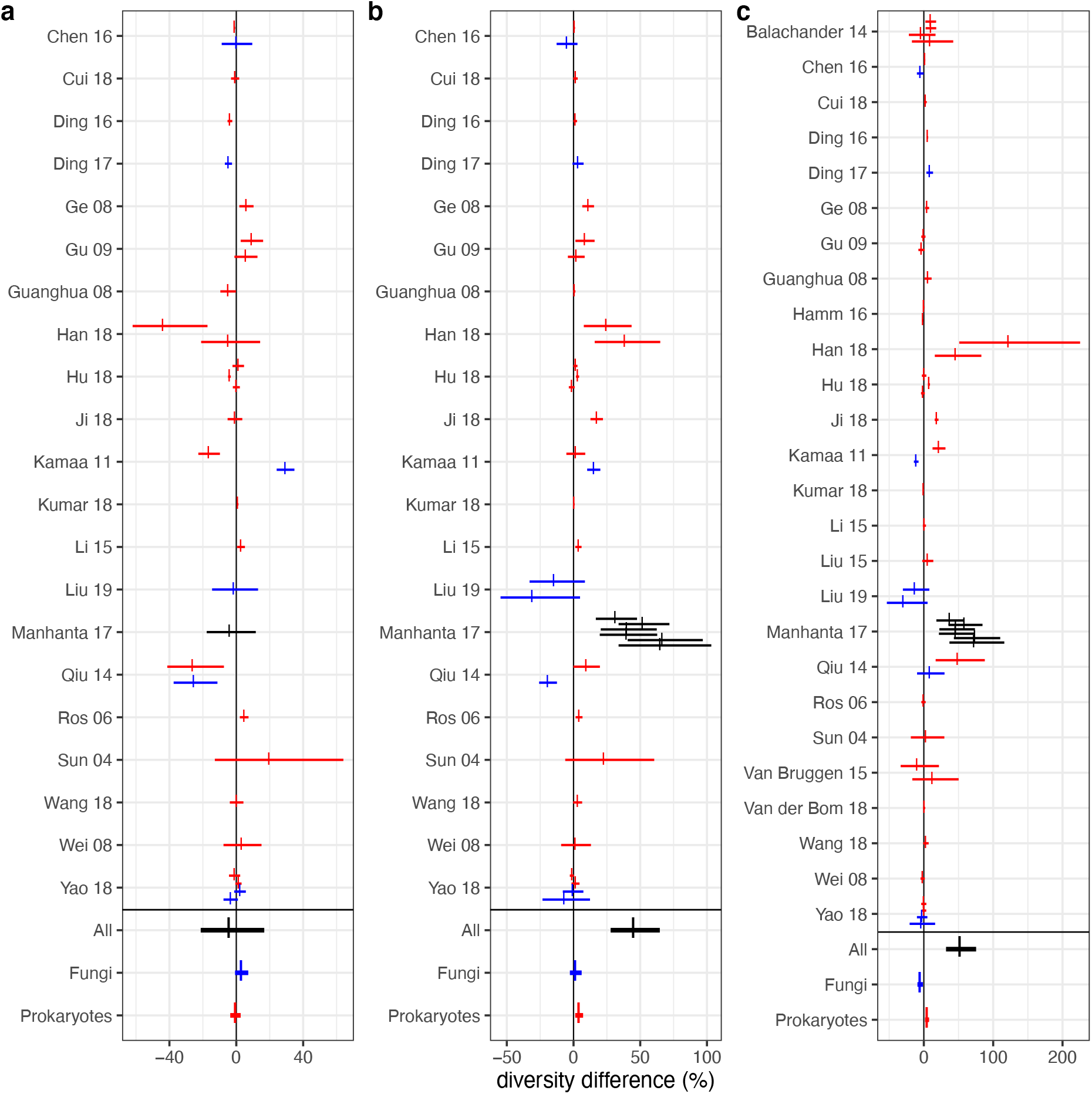
Taxonomic diversity effect sizes. a) NPK vs. CON, b) ORG vs. CON, c) ORG vs. NPK. Horizontal bars and ticks show 95% confidence intervals and means for effect sizes from the log-transformed ratio of means, back-transformed to give percentage differences. Summary estimates are given at the bottom of each plot, for fungi (blue) and prokaryotes (red). Random intercepts were fitted per Study.

## DISCUSSION

We found significant, but varying, effects of both NPK and organic fertilizers on soil microbe taxonomic and functional diversity. Taxonomic diversity in organic treatments compared with NPK was greater for prokaryotes but not significantly different for fungi. We found no significant difference in taxonomic diversity between NPK and control. Functional diversity was significantly greater in NPK compared with control, and in organic treatment compared with NPK. However, the diversity differences we found were small, indicating that choice of fertilization has marginal effects on this measure of soil microbial community structure. A quantitative review on the effects of fertilizers on soil fungal diversity found that mineral fertilizers tended to reduce diversity while organic fertilizers had no significant effect (Ye et al., 2020), although the analysis did not consider sample sizes or variances of study data and so the findings are difficult to interpret.

A large meta-analysis of the impacts of global change factors found that N addition alone significantly reduced soil bacterial but not fungal Shannon diversity, particularly in agricultural systems (Zhou et al., 2020). The same meta-analysis did not detect significant effects of NPK overall, but the effects of organic fertilizers were not investigated. The most important moderator was found to be pH, which strongly controlled the response ratio of diversity in response to global change factors. In line with our findings, N and NPK treatments were found to reduce pH. In contrast, we found a buffering effect of manure fertilizers on soil pH, as reported in earlier studies (reviewed in Köninger et al., 2021). This buffering effect, along with nutrient and organic matter content, are considered the main benefits of manure fertilizers for soil biodiversity (Köninger et al., 2021).

Several factors must be considered when interpreting the effects of fertilizers on soil microbial functional and taxonomic diversity (Fig. 4). First, while in many studies the amount of nitrogen applied in organic and NPK treatments was similar, farmyard manure is far more physically, chemically and biologically complex. In addition to NPK, manures contain undigested plant matter (lignin, cellulose, hemicellulose), lipids, carbohydrates, proteins, and nutritive elements (e.g. magnesium, iron, manganese, zinc, copper) (Levi-Minzi et al., 1986). If structural complexity and heterogeneity are enhanced by the addition of organic matter across spatial scales (Lehmann et al., 2008), this could in itself increase microbial diversity through provision of ecospace (Vos et al., 2013). Provision of additional energy (organic carbon) and micronutrients in manures could sustain a greater diversity of microbes via the species-energy hypothesis which predicts that more species can be sustained in ecosystems supporting more individuals (Clarke and Gaston, 2006), since manure increases soil microbial biomass more than NPK (Lori et al., 2017). However, one potential risk of manure fertilizers is that high concentrations of heavy metals such as zinc and copper can lead to soil accumulation and reductions in microbial diversity (Köninger et al., 2021).

**Fig. 4.**
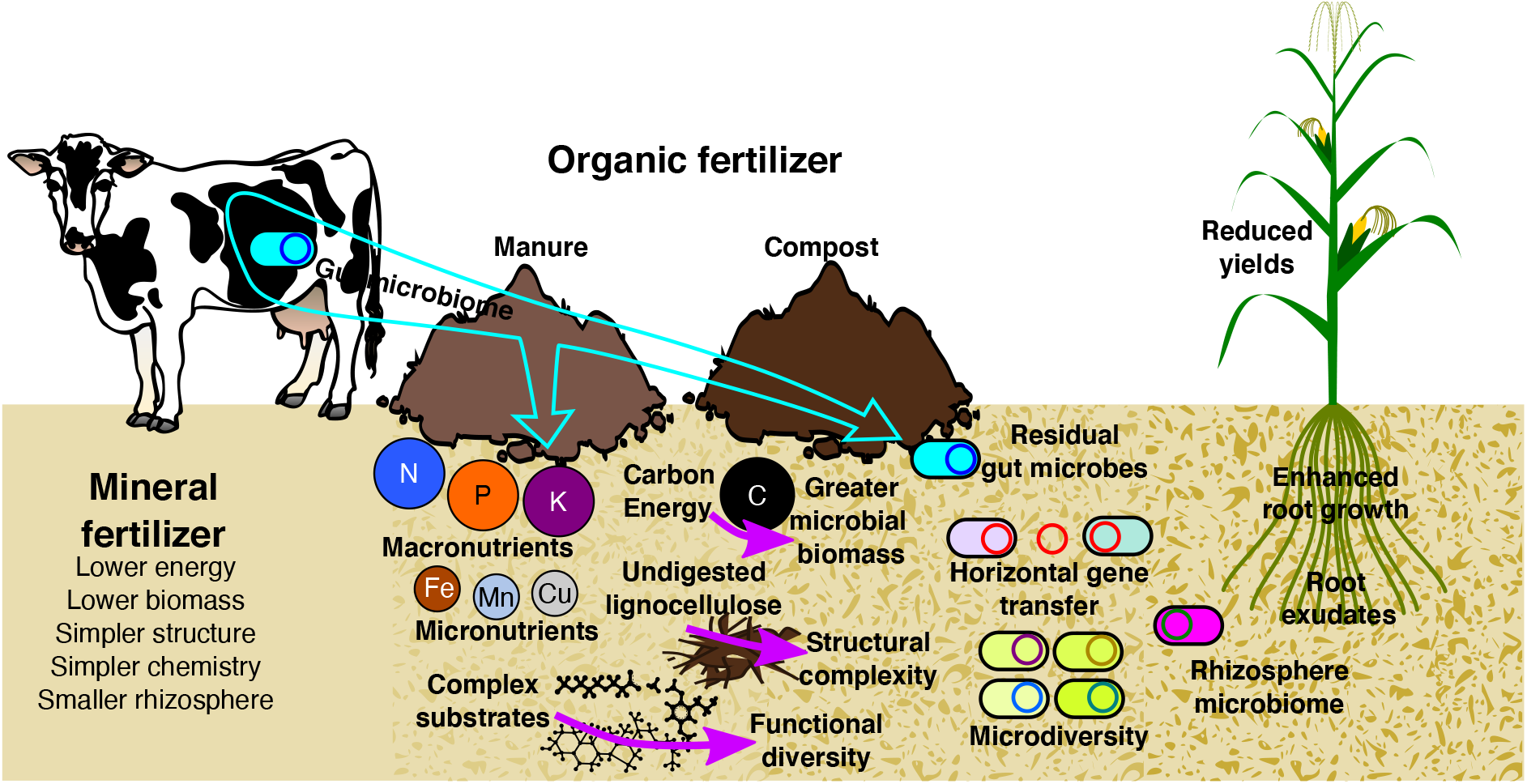
Representation of processes potentially influencing fertilizer affects on soil microbial diversity. See Discussion for details. Graphical elements from https://svg-clipart.com/ and http://clipart-library.com/

Second, microbial communities may be indirectly influenced by changes in crop plants. While organic agriculture tends to be less productive in terms of crop yields (de Ponti et al., 2012; Hijbeek et al., 2017), there is some evidence that root development is greater in organic farming (T. Hu et al., 2018). Plant rhizospheres tend to harbour the greatest microbial diversity of all terrestrial ecosystems (Thompson et al., 2017), hence it is possible that enhanced root development and exudation of organic compounds influence the soil diversity measured in these studies. Third, there is the gut microbial community residing within the manure itself. Both fresh and composted manures have high fungal and prokaryote diversity, with community composition changing as composting proceeds (Meng et al., 2019). Composting can reduce the presence of undesirable microbes in manure, for example those carrying antimicrobial resistance genes (Gou et al., 2018), but gut microbes (including human pathogens) can survive for long periods in compost-amended soils (Sharma and Reynnells, 2016). Hence, increased soil microbial diversity in organic systems may be due to persistence of gut microbes.

Few studies reported both taxonomic and functional diversity, hence we were unable to determine a relationship between these two metrics. Microbial functional diversity tends to be considered from the perspective of potential functions inferred from gene sequences (Escalas et al., 2019). In contrast, we report results from studies of the diversity of actual functions (metabolization of carbon sources) carried out by the soil microbial community. The relationship between soil taxonomic diversity and ecosystem function (as opposed to functional diversity) tends to be positive (Bardgett and Putten, 2014; Maron et al., 2018; Philippot et al., 2013). Soil ecosystem multifunctionality (the capacity of soils to sustain many functions simultaneously) increases with microbial diversity (Delgado-Baquerizo et al., 2016), while taxonomic and functional gene diversity are closely correlated (Zhang et al., 2019). However, taxonomic and functional diversity can be decoupled by the process of horizontal gene transfer, because different taxa can perform similar tasks through shared genes (Zhang et al., 2019). In addition, the presence of physiologically-distinct subgroups within OTUs commonly defined by > 97 % sequence similarity (so-called ‘microdiversity’) means that functional diversity can be large in groups of apparently identical microbial taxa (Larkin and Martiny, 2017).

Organic farming has been promoted as a more sustainable alternative to conventional agriculture, because of greater energy efficiency, potentially closed nutrient cycles and increased biodiversity (Reganold and Wachter, 2016). These potential benefits come at a cost of reduced productivity compared with convential farming (de Ponti et al., 2012; Hijbeek et al., 2017), though there is some evidence that organic production can eventually catch up with conventional yields and provides greater spatial and temporal stability (Schrama et al., 2018). Environmental impacts of organic agriculture include greater land use and eutrophication potential per unit of food produced, contradicting the aim of closed nutrient cycles (Clark and Tilman, 2017). Our results suggest that, when considering organic crop fertilization alone and as a strict alternative to NPK (rather than a mixed system of manure-derived and NPK fertilizer), microbial functional diversity and bacterial taxonomic diversity are slightly, but significantly greater in organic systems while the effect on fungal taxonomic diversity is unclear. Despite the increases in microbial diversity under organic vs. conventional fertilization we found, organic farming systems still often exhibit higher eutrophication potential than conventional systems due to temporal mismatching between fertilizer addition and plant demand (Clark and Tilman, 2017). Therefore, it appears that the diversity of the microbial community is likely not the most important factor governing rates of nutrient loss from organic vs. conventional systems. While we did not compare combined treatments (e.g. NPK with manure or compost) with single treatments, farmers commonly apply a diversity of fertilizers. For example, a comparison of low-input organic systems with conventional mixed (NPK and manure) and conventional NPK-fertilized systems in Switzerland found the greatest bacterial α-diversity in the organic systems, followed by the mixed and NPK-only systems (Hartmann et al., 2015). Some studies which were not included in the meta-analysis employed manure-free organic treatments, finding no effect (Wu et al., 2015) or variable effects (Wang et al., 2016) on soil microbial diversity.

Many studies addressing soil microbial diversity in response to fertilization could not be included in our meta-analysis because of incomplete or unclear reporting. Without giving specific examples, we found that many studies did not provide uncertainties (standard error or standard deviation) for parameter estimates, gave unclear descriptions of the fertilisation routines and inputs, or reported alternative measures of diversity. Of the studies we included, several did not report basic soil chemistry metrics such as pH and nitrogen content. Given the importance of pH in determining soil microbial diversity (Zhou et al., 2020), and the central relevance of nitrogen in these experiments, any future research should report these variables at minimum. We were only able to include a relatively small number of studies in our meta-analysis, though we note that in the Cochrane Library of medical meta-analyses, the median number of studies is seven or below (von Hippel, 2015). The Cochrane Collaboration has been instrumental in developing methods of systematic review and meta-analyses for evidence-based medicine(Higgins et al., 2011). One reason for the lack of experimental comparisons of organic and mineral fertilizer treatments could be that organic agriculture comprises only a tiny fraction, around 1 per cent by area, of agriculture globally (Meemken and Qaim, 2018). We found very large residual heterogeneity among effect sizes, i.e. most of the variation in effect sizes among studies remains unexplained. Further research will reveal whether the mean effects we detected are general, and what other factors help explain variation among soils, climates, locations and experimental treatments. To achieve this, it is critical that detailed and complete data on key variables are reported. In addition, better understanding of soil ecosystem functioning will be achieved via analysis of the taxa and functional genes identified within samples, in addition to summary metrics like diversity (Hartmann et al., 2015). In this way, the powerful new bioinformatics tools at our disposal can be harnessed to fully understand the relationships between microbial communities and soil health in agricultural systems.

## DECLARATIONS

### AVAILABILITY OF DATA

The data used in the meta-analysis will be made available on publication of the manuscript.

### CODE AVAILABILITY

Code used in the meta-analysis is available from the author by reasonable request.

### AUTHORS CONTRIBUTIONS

DB developed the research. DB and VR collected data, analysed results and wrote the paper.

### ETHICS APPROVAL

Not applicable

### CONSENT TO PARTICIPATE

Not applicable

### CONSENT FOR PUBLICATION

Not applicable

## SUPPLEMENTARY MATERIAL

### SUPPLEMENTARY FIGURES

**Fig. S1.**
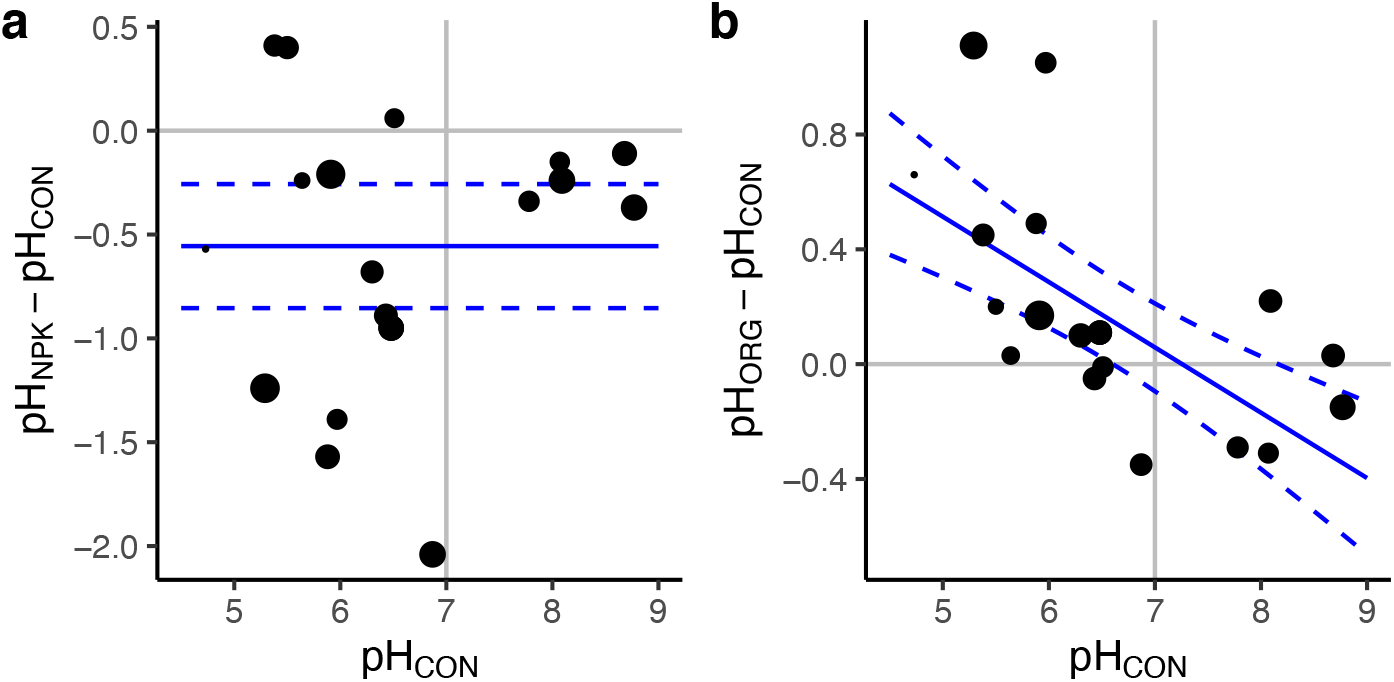
Effect of fertilization on soil pH. a) Mean difference between NPK and CON vs. mean CON pH. Blue solid and dashed lines show mean difference (−0.59 ± 0.19 SE, Z = -3.09, p = 0.002) and 95% CI (−0.97, -0.22) of meta-analysis fit. b) Mean difference between ORG and CON vs. mean CON pH. Blue solid and dashed lines show mean and 95% CI of meta-regression fit. Slope mean -0.27 ± 0.087, Z = -3.08, p = 0.002, 95% CI = -0.44, -0.10. Points show means from individual studies with size proportional to log(1/variance).

**Fig. S2.**
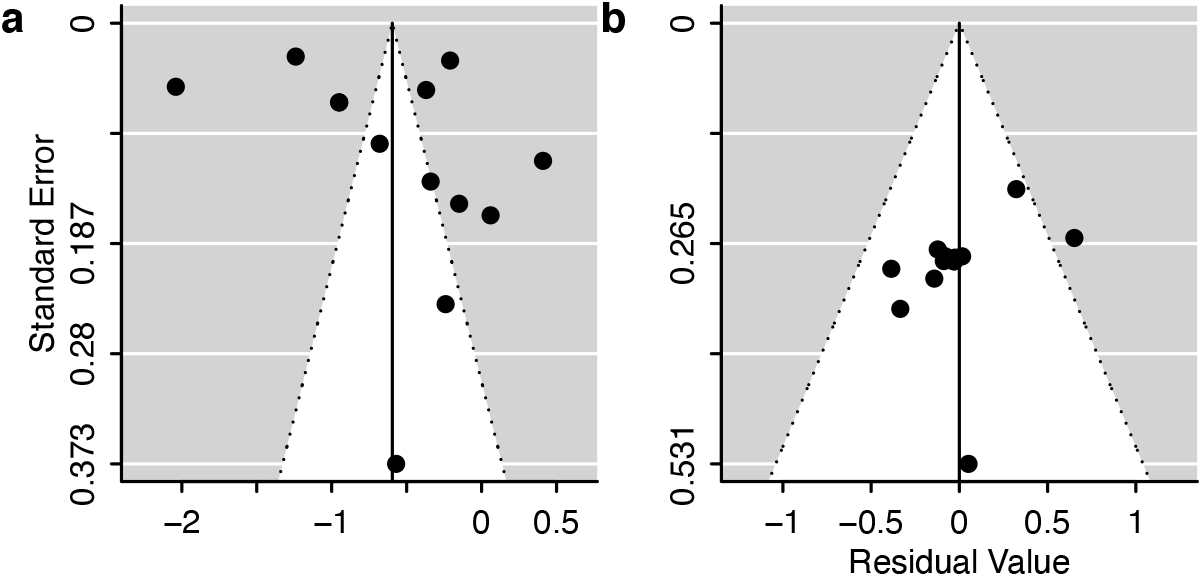
Funnel plots for soil pH meta-analysis. Kendall’s τ and p-values are given for rank correlation tests for funnel plot asymmetry. a) NPK vs. CON (τ = 0.14, p = 0.50), b) ORG vs. CON (τ = 0.25, p = 0.24).

**Fig. S3.**
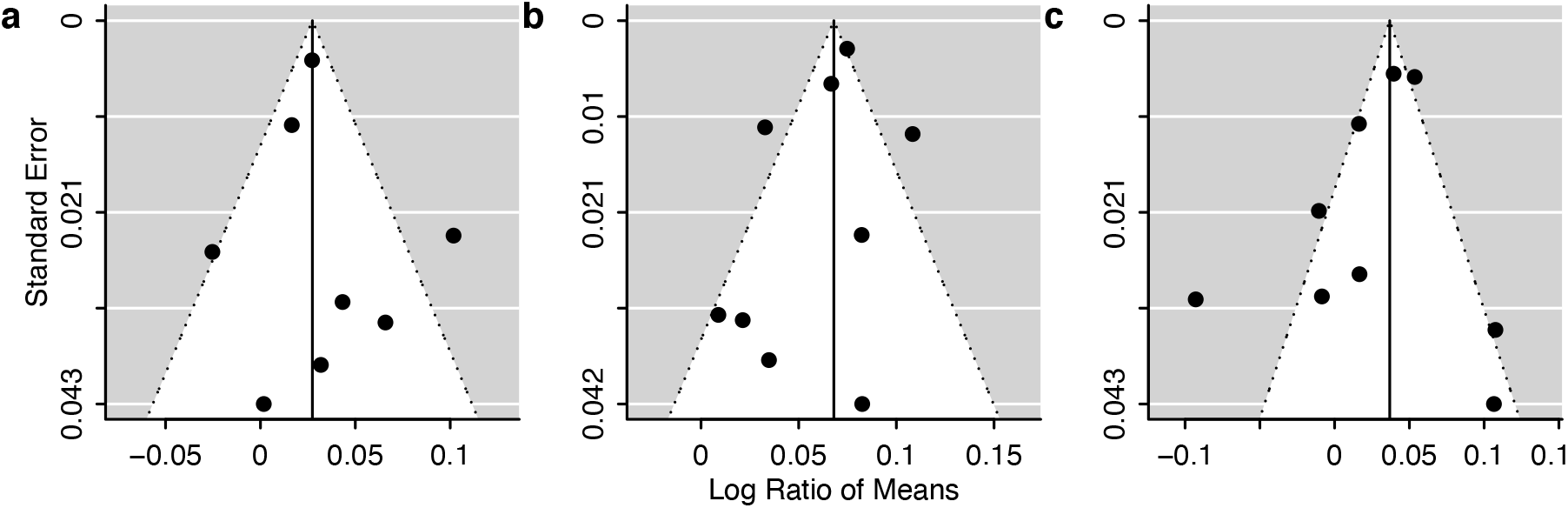
Funnel plots for functional diversity meta-analysis. Kendall’s τ and p-values are given for rank correlation tests for funnel plot asymmetry. a) NPK vs. CON (τ = 0.00, p = 1.00), b) ORG vs. CON (τ = -0.11, p = 0.76), c) ORG vs. NPK (τ = -0.16, p = 0.91).

**Fig. S4.**
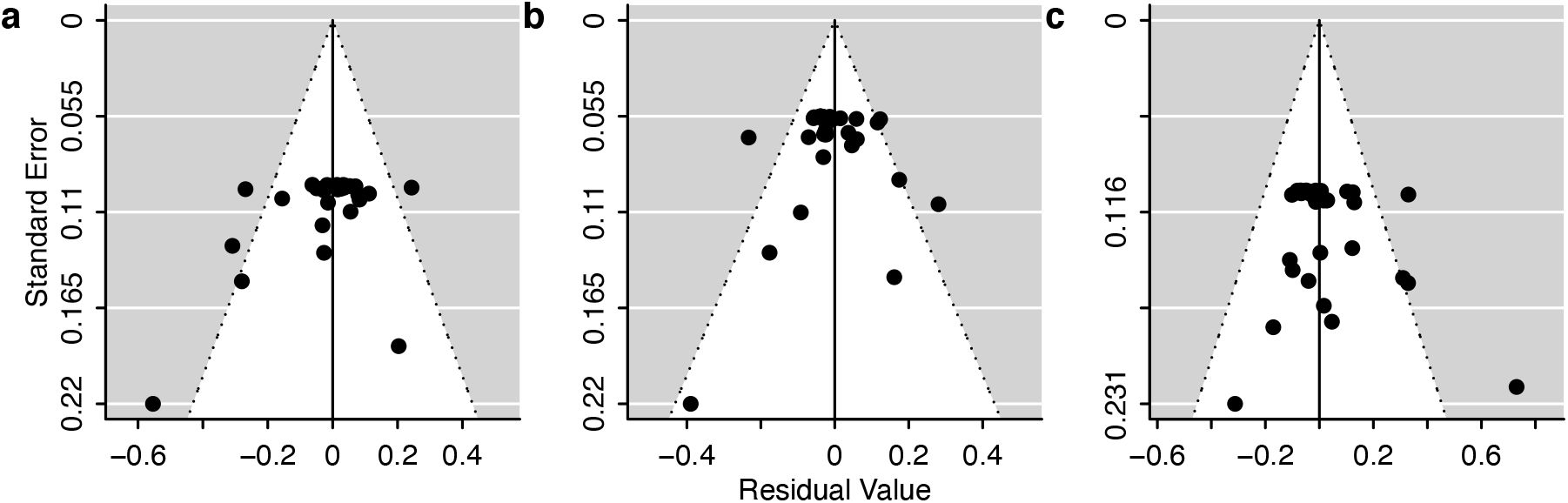
Funnel plots for taxonomic diversity meta-analysis. Kendall’s τ and p-values are given for rank correlation tests for funnel plot asymmetry. a) NPK vs. CON (τ = -0.10, p = 0.44), b) ORG vs. CON (τ = 0.02, p = 0.91), c) ORG vs. NPK (τ = 0.12, p = 0.26).

### SUPPLEMENTARY TABLES

**Table S1.**
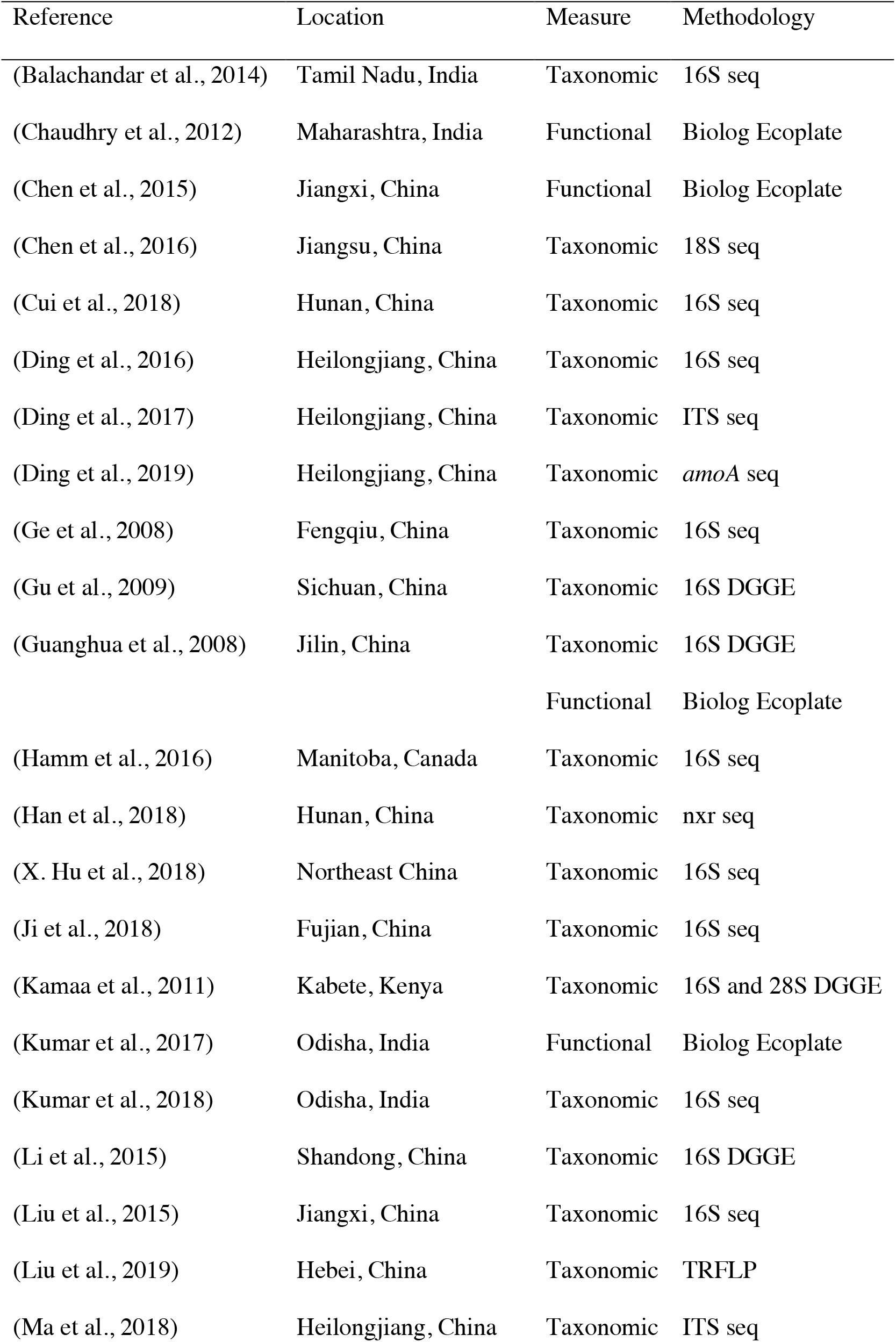
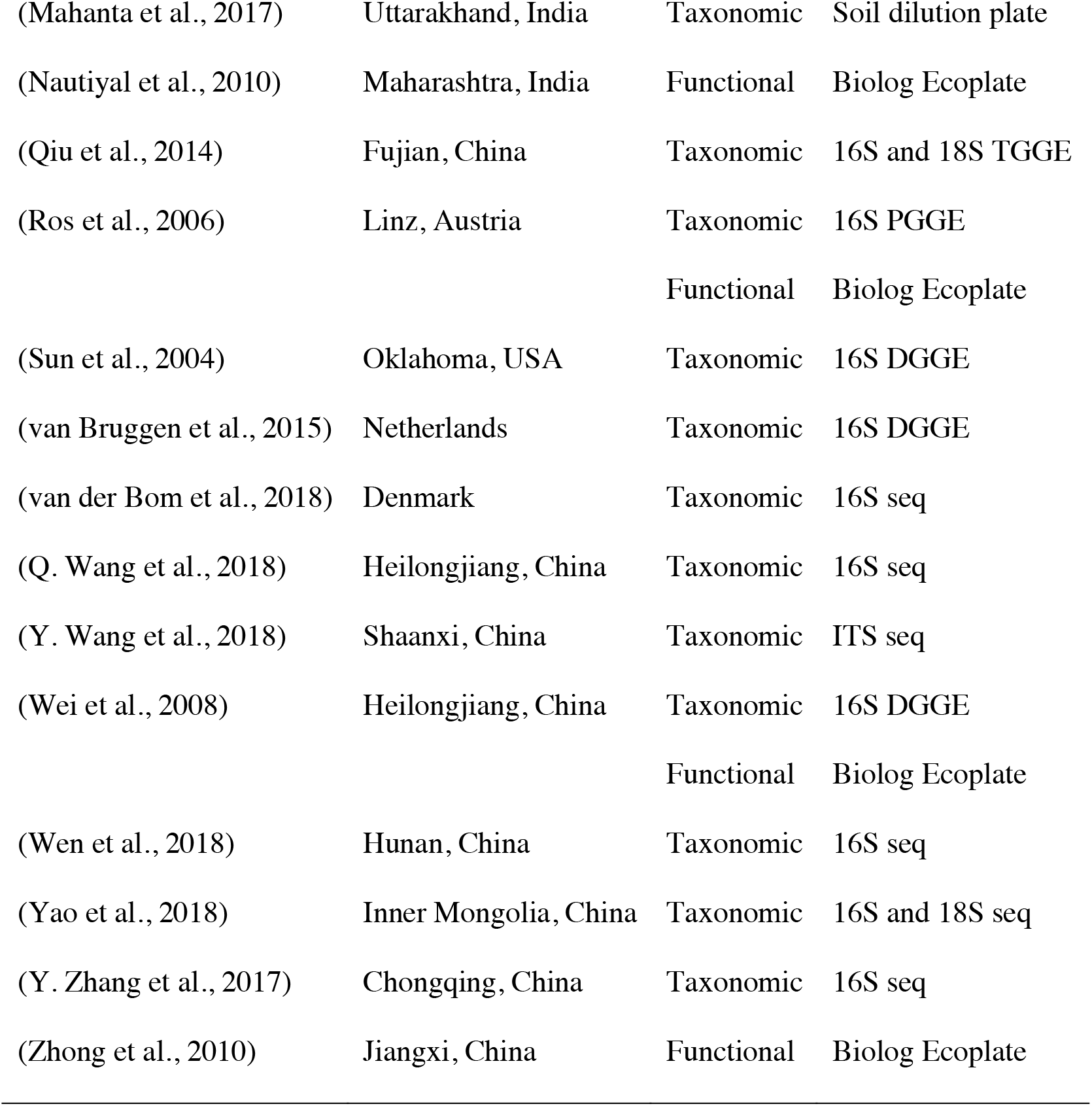
Summary of studies included in the meta-analysis (alphabetical order).

**Table S2.** Meta-analysis diagnostics.

| Test | I<sup>2</sup> (%) | Q | df | P |
| --- | --- | --- | --- | --- |
| H<sub>fun</sub> NPK vs. CON | 73.5 | 16.5 | 7 | 0.0210 |
| H<sub>fun</sub> ORG vs. CON | 84.6 | 29.1 | 8 | 0.0003 |
| H<sub>fun</sub> ORG vs. NPK | 82.3 | 42.3 | 8 | <0.0001 |
| H<sub>tax</sub> NPK vs. CON | 96.6 | 570.0 | 29 | <0.0001 |
| H<sub>tax</sub> ORG vs. CON | 91.4 | 256.6 | 30 | <0.0001 |
| H<sub>tax</sub> ORG vs. NPK | 96.4 | 874.6 | 40 | <0.0001 |
| pH<sub>NPK</sub> - pH<sub>CON</sub> vs. pH<sub>CON</sub> | 99.0 | 1353.5 | 11 | <0.0001 |
| pH<sub>ORG</sub> - pH<sub>CON</sub> vs. pH<sub>CON</sub> | 94.5 | 10.9 | 10 | 0.0009 |

